# Whole Genome Selective Sweeps Analysis in Pakistani Kamori Goat

**DOI:** 10.1101/2021.01.25.428182

**Authors:** Rashid Saif, Jan Henkel, Tania Mahmood, Aniqa Ejaz, Saeeda Zia

## Abstract

Natural and artificial selection fix certain genomic regions of reduce heterozygosity which is an initial process in breed development. Primary goal of the current study is to identify these genomic selection signatures under positive selection and harbor genes in Pakistani Kamori goat breed. High throughput whole genome pooled-seq of Kamori (n = 12) and Bezoar (n = 8) was carried out. Raw fastq files were undergone quality checks, trimming and mapping process against ARS1 reference followed by calling variant allele frequencies. Selection sweeps were identified by applying pooled heterozygosity (*Hp*) and Tajima’s D (TD) on Kamori while regions under divergent selection between Kamori & Bezoar were observed by Fixation Index (F_ST_) analysis. Genome sequencing yielded 619,031,812 reads of which, 616,624,284 were successfully mapped. Total 98,574 autosomal selection signals were detected; 32,838 from *Hp* and 32,868 from each F_ST_ & TD statistics. Annotation of the regions with threshold (−*ZHp* ≥ 5, TD ≤ −2.72 & F_ST_ ≤ 0.09) detected 60 candidate genes. The top hits harbor Chr.1, 6, 8 & 21 having genes associated with body weight (*GLIS3, ASTE1*), coat color (*DOCK8*, *MIPOL1*) & body height (*SLC25A21*). Other significant windows harbor milk production, wool production, immunity, adaptation and reproduction trait related genes. Current finding highlighted the under-selection genomic regions of Kamori breed and likely to be associated with its vested traits and further useful in breed improvement, and may be also propagated to other undefined goat breeds by adopting targeted breeding policies to improve the genetic potential of this valued species.

## 1 Introduction

Domestic goat breeds have been under well documented and strong selection for several years through natural and artificial methods for various quantitative traits especially meat, milk and reproduction. The selection strategies impose selection pressure on particular regions of the genome that control such traits. For instance, intensive crossbreeding strategy was applied on Pakistani breeds, Pak-Angora and Beetal with Hairy goat, for bulk quality production of Mohair [1]. Similarly, hybrid Kamori goat (Kamori x Patairee) are popular in Pakistan for beautiful coat color, appearance and being less expensive than its pure breed [2]. So ascertaining the genes and genomic regions under positive selection affected due to selection force is vital for understanding the phenotypic diversity among breeds due to genetic variations [3]. Whole genome analysis based on population genetic statistics have been developed that significantly asses the selective sweeps in all livestock species without known phenotypes. These include; F_ST_ for genomic differentiation [4], cross-population composite likelihood ratio (XP-CLR) that uses variability in allele frequencies at linked loci only between two populations, *Hp* uses variability estimator in allele counts [5], extended haplotype homozygosity (EHH) approach using SNP data [6], TD statistics to calculate possible distortions in distribution of allele frequencies [7] and others have been used to reveal selection footprints in various livestock species.

In this study, we used three statistical methods (i) *Hp* (ii) TD and (iii) F_ST_ to explore genomic selective sweeps in Kamori goat vs. the wild ancestor Bezoar from Switzerland. Kamori is famous for its long ears, neck and unique coat colors usually dark brown with small dark or coffee colored patches over its entire body. They are medium to large size milch breed mainly found in Nawabshah, Dadu and Larkana districts in Sindh province of Pakistan. On average a Kamori doe gives ~1.5 liters milk [1]. The primary goal of this research was to identify genomic regions and underlying candidate genes affected by selection in Kamori goat.

## 2. Materials and methods

### 2.1 Animals and high-depth WG pooled sequencing

Genomic DNA was extracted for high throughput sequencing, from Kamori (n = 12) using TIANGEN biotech (Beijing) CO., LTD and extraction from Bezoar goat (n = 8) was carried out at Institute of Genetics, University of Bern, Switzerland from whole blood samples. Same amount of extracted DNA per individual was mixed into a single pool. Whole genome sequencing was performed using Illumina HiSeq 2000 platform that generated 150bp paired-end ~300mio reads. Sequencing files are available at European Nucleotide Archive (ENA) under Project ID: PRJEB23815. Summary of the characteristics (Table 1) along with the figures of all three breeds used in this study are shown in Fig. (1).

**Table 1.**
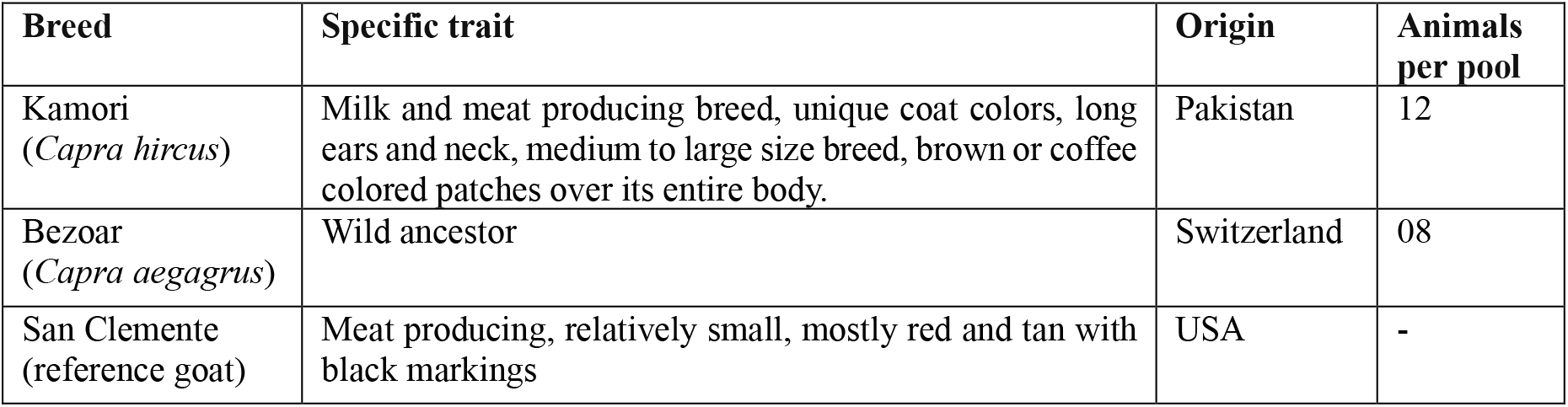
Features summarizing Kamori (breed), Bezoar (wild ancestor) and San Clemente (reference goat) used in this study.

**Fig. (1).**
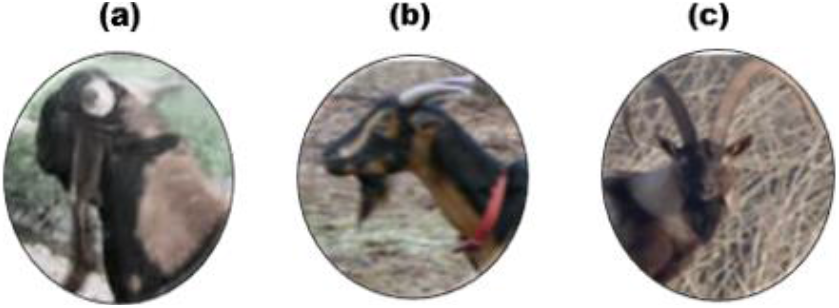
Representative animals. (a): Kamori (subject goat). (b): San Clemente (reference goat). (c): Bezoar goat (wild ancestor).

### 2.2 Calling SNVs from whole genome pooled-seq data

Filtration of both pools fastq files for base quality was done by Trimmomatic (v0.36) using SLIDINGWINDOW:4:20 MINLEN:2 parameters. Filtered reads were aligned to ARS1 reference genome assembly using BWA-MEM algorithm v0.7.17 [8] followed by SAM to BAM conversion using its samtools view feature. Sorting on coordinate basis and tagging adapter sequences was accomplished using SortSam and MarkDuplicates features of Picard tools [9]. Quality of reads were checked using FastQC (v0.11.8) software. Samtools mpileup was run on both pools bam files together and on single Kamori bam file to call SNVs. Resulting mpileup and pileup files were subjected to Popoolation2 v1.201 tool [10], mpileup2sync.jar and snp-frequency-diff.pl scripts, that generated synchronized (sync) combined mpileup and separate sync pileup file.

### 2.3 Selective sweep detection using Hp, TD and F_ST_ statistics

An in-house Ruby script with window-size of 150kb was applied to calculate −*ZHp* scores by first applying *Hp* = 2Σn_MAJ_Σn_MIN_/(Σn_MAJ_+Σn_MIN_)^2^ and then Z-transforming it by using −*ZHp*= -(*Hp*-μ*Hp*/σ*Hp*) to search for positive selection hits [5]. On separate Kamori pileup file, Popoolation v1.2.2 tool’s script, variance-sliding.pl was run [7] to compute classical Tajima’s D applying D_b_._pool_ = d_b.pool_/√Var(d_b.pool_). For genomic differentiation between the two goat breeds, Popoolation2 v1.201 tool’s script, fst-sliding.pl based on F_ST_ = s^2^/p̄(1-p̄) + s^2^/r [4] was applied on each SNV value obtained in combined sync file with 50% overlapping window.

### 2.4 Data visualization

Manhattan plots of – *ZHp*, TD and F_ST_ values were constructed using manhattan function of qqman package while SNP density graph of these scores was constructed using CMplot package on R software [11]. In addition, the – *ZHp*, TD and F_ST_ scores were constructed against the theoretical quantiles as Q-Q plots using qqnorm function while the distribution of these values across the no. of windows was plotted as histogram using hist function on R [12].

## 3. Results

### 3.1 Quality control and pooled-seq data processing

Genome sequencing yielded 619,031,812 total reads which are further trimmed. Almost 616,624,284 reads with a coverage of 99.61% are mapped against the ARS1 reference genome assembly which are quality checked for further onward selection signature study Fig. (S1). SNV calling generated 98,574 autosomal selection signals including 32,838 from *Hp* and 32,868 variants each from F_ST_ and TD statistics. The distribution of number of SNPs within 100 MB window size by each of the method is shown in Fig. (S2). Likewise, the Q-Q plot of – *ZHp*, TD and F_ST_ values across all autosomes are observed in black line against the expected standard normal distribution in red line along with the frequency distribution graphs of – *ZHp*, TD and F_ST_ values in Fig. (S3; S4 and S5).

### 3.2 Candidate selection signals and harbor genes

The windows under positive selection in Kamori goat obtained after setting thresholds (−*ZHp* ≥ 5, TD ≤ −2.72 and F_ST_ ≤ 0.09) on the basis of previously published data and rationale observation of our own data, are further fine-mapped and annotated which revealed genes related to body mass, coat color, wool type, milk production, immunity, adaptation to environment, body height and reproduction (Table 2).

**Table 2.** Genes under selection in Kamori goat revealed by *Hp*, F_ST_, TD and its associated traits.

| Under selection genes | Related traits | References |
| --- | --- | --- |
| *USP33, HMBOX1, GLIS3, TJP2, FAM189A2, APBA1, MYCBP2, KLHL1, RAPH1, STIM1, ASTE1, ATP8B3, RAPGEF6, SDHAF4, B3GNT6, FAM168A, ACSF3* | Meat production, body mass and body weight | [13-15] |
| *ANKRD11, RERE, PRP6* | Reproduction and embryonic development | [16] |
| *DMRT2, DMRT1, ZFPM1, ATP2C1* | Milk protein composition and production | [17, 18] |
| *SAXO1, ADAMTSL1, DUSP15, EIF2S2, PIEZO1, CDH15, CPNE7, TDP2, SPIRE2, RIC1, TPX2, STAG1, CCDC91, ZNF609, MGA, ATAD2B, SIK3, SLC255A21, TET1, ADAMTS6, NCAPG* | Body height | [9, 19, 20] |
| *FH, WDPCP, TRMT11* | Environmental adaptation | [21] |
| *SLC38A8* | Silky hairs/wool type | [22] |
| *KIT, KANK, DEF8, DOCK8, MLANA, RALY, MIPOL1, IRF4* | Coat color | [10, 23] |
| *CBFA2T3, RFX3, CTLA4* | Immunity | [24] |

### 3.3 Signatures of positive selection by Hp analysis

The *Hp* analysis of Kamori breed returned 62 windows under selection with – *ZHp* ≥ 5 Fig. (2). The top hits appear on Chr.8:40,500-40,650 kb, Chr.6:70,800-70,950 kb and Chr.8:40,425-40,575 kb regions having – *ZHp* scores 5.96, 5.91 and 5.90 containing 321, 280 and 267 number of SNPs respectively. The 62 putative regions under selection harbor 28 known genes related to body weight, body height, coat color, reproduction and milk protein composition while 10 windows have LOC genes and 11 regions have no genes (Table S1).

**Fig. (2).**
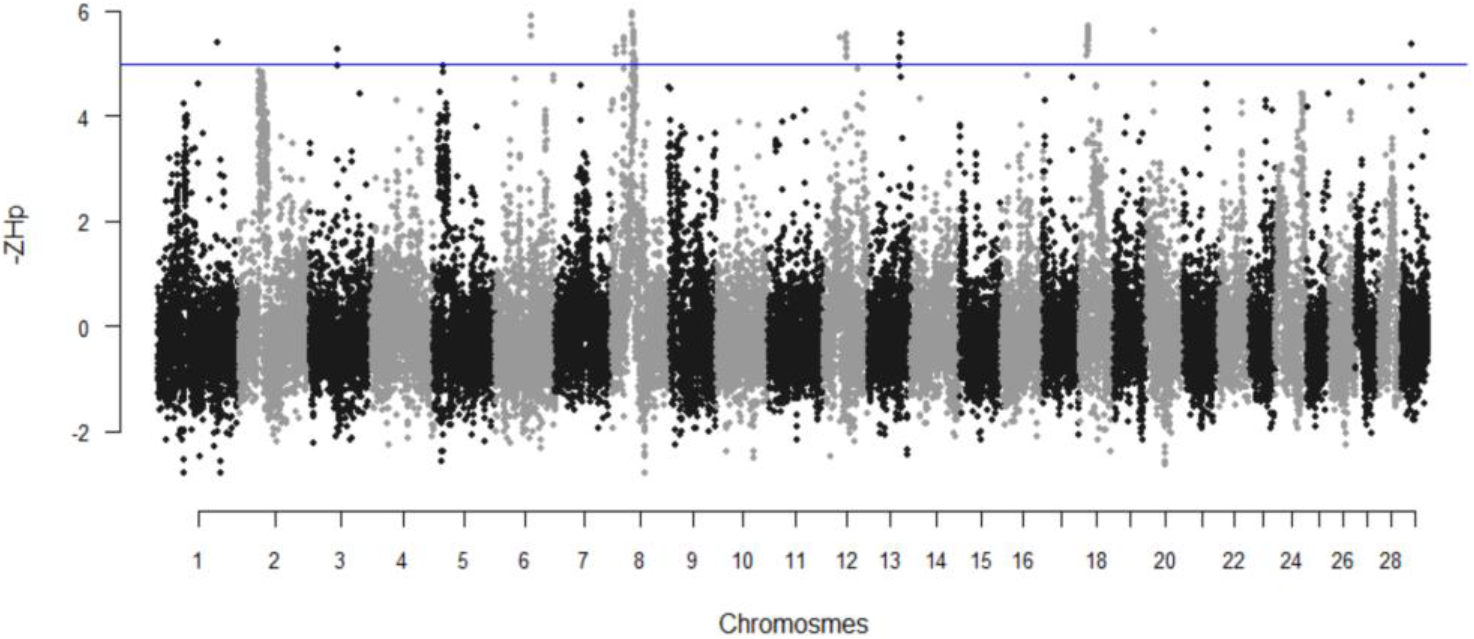
Manhattan plot of – *ZHp* analysis on Kamori goat breed. The blue horizontal line directs the suggested significant cutoff threshold of – *ZHp* ≥ 5 for better hits. The sliding window approach was used with window size of 150 kb and 75 kb step size considering all autosomes.

### 3.4 Regions under purifying selection by Tajima’s D statistics

Based on the calculations of TD, 63 significant windows (having TD ≤ −2.72) corresponding to the regions that have undergone a recent bottleneck or a selective sweep are revealed Fig. (3). The top most significant windows with TD values −2.81 and −2.79 are located on Chr.8:4,025-4,175 kb and 40,500-40,650 kb region possessing 2,218 and 2,188 SNPs respectively. Twenty three genes are identified lying on these putative selective sweeps that are associated with bone length/bone height, body weight/meat production, milk protein, immunity, coat color and cold-induced thermogenesis for adaptation to the environment. Twelve of the regions have LOC genes while 07 putative windows are devoid of genes (Table S2).

**Fig. (3).**
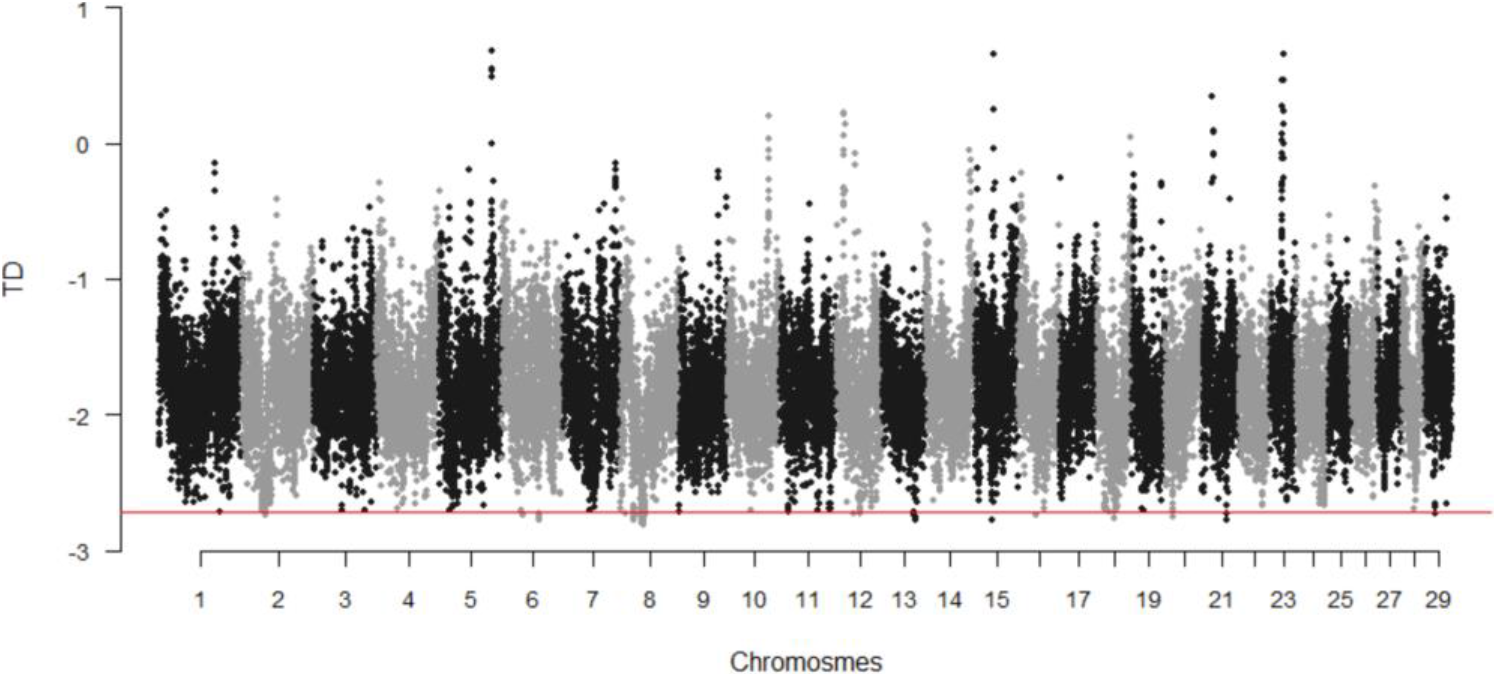
Manhattan plot showing TD scores from Kamori goat breed. The red horizontal line indicates the stringent threshold of TD ≤ −2.72 that is computed using 50% overlapping window size such that each dot represents a 150kb window.

**Fig. (4).**
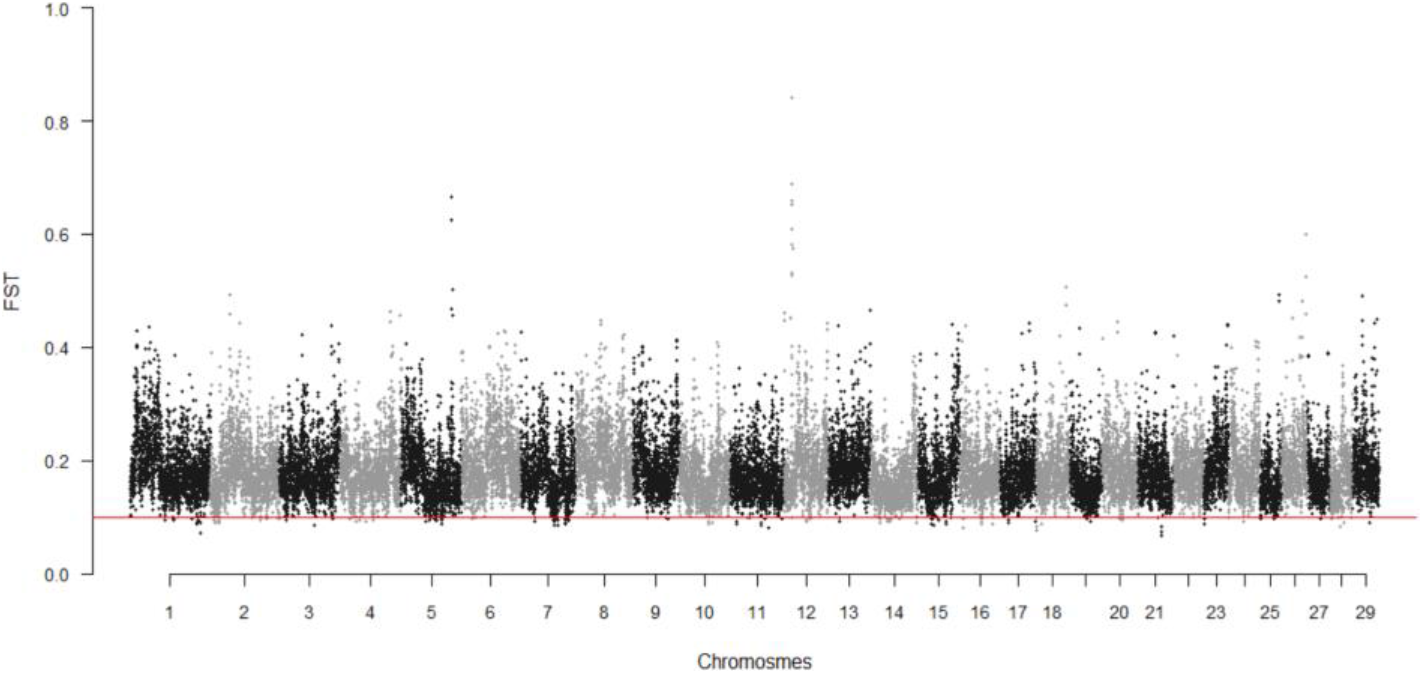
Manhattan plot based on F_ST_ values distributed across all autosomes of Kamori goat. The preferred threshold of F_ST_ ≤ 0.09 is illustrated by horizontal red line. By using sliding window approach, window size was set to 150 kb and step size 75 kb.

### 3.5 Regions under divergent selection by F_ST_ analysis

F_ST_ methodology is applied on Kamori and Bezoar to focus on the genomic regions that differ between the two. Thirty four candidate regions are divulged after adjusting the F_ST_ threshold to ≤ 0.09. Of these significant windows, highly differentiated regions are positioned on Chr.21:46,875-47,025 kb, 46,800-46,950 kb and on Chr.1:138,450-138,600 kb region having 0.66, 0.73 and 0.72 F_ST_ values comprising 3,995, 4,081 and 4,919 SNPs respectively. Fine mapping of regions under selection revealed 23 genes related to milk protein and fat, body height and weight, immunity, reproduction, coat color, adaptation to high altitude and in cold environment and silky hair, 7 LOC genes and only 1 window has no gene (Table S3).

### 3.6 Congruent Selection signatures identified by – ZHp, TD and F_ST_ approaches

Table 3 list the putative selection signature and underlying genes that are observed above the preferred thresholds by at least two of the three methods. Interestingly, 23 windows that harbor 15 genes are commonly selected by – ZHp and TD while none of the signals by both methods corresponds to the signals generated by F_ST_.

**Table 3.** Common selective sweeps and genes identified by three approaches– *ZHp*, *TD and F_ST_.*

| Chr. | Common selective sweeps | Statistical methods | Gene | Selected traits | Significant values |
| --- | --- | --- | --- | --- | --- |
| 6 | 70,725-70,875 kb | *Hp*, TD | *KIT* | Coat color | -ZHp = 5.532, TD = -2.734 |
| 6 | 70,800-70,950 kb<br>70,875-71,025 kb | *Hp*, TD | *LOC102174549* | - | -ZHp = 5.91, 5.72,<br>TD = -2.75, -2.77 |
| 8 | 9,450-9,600 kb<br>9,525-9,675 kb | *Hp*, TD | *HMBOX1* | Body weight,<br>meat production | -ZHp = 5.30, 5.18,<br>TD = -2.72, -2.73 |
| 8 | 25,200-25,350 kb<br>25,275-25,425 kb<br>25,350-25,500 kb<br>25,425-25,575 kb | *Hp*, TD | *ADAMTSL1* | Body height | -ZHp = 5.22, 5.44, 5.33,<br>5.48 TD = -2.74, -2.75,<br>-2.74, -2.77 |
| 8 | 38,850-39,000 kb | *Hp*, TD | *RIC1* | Body height | -ZHp = 5.18, TD = -2.76 |
| 8 | 40,350-40,500 kb | *Hp*, TD | *TRNAG-UCC* | tRNA gene | -ZHp = 5.73, TD = -2.78 |
| 8 | 40,425-40,575 kb<br>40,500-40,650 kb | *Hp*, TD | *GLIS3* | Body weight | -ZHp = 5.90, 5.96,<br>TD = -2.78, -2.79 |
| 8 | 43,200-43,350 kb<br>43,275-43,425 kb | *Hp*, TD | *DMRT2* | Milk protein | -ZHp = 5.44, 5.54<br>TD = -2.76, -2.77 |
| 8 | 43,350-43,500 kb<br>43,425-43,575 kb | *Hp*, TD | *DMRT1* | Milk protein | -ZHp = 5.53, 5.18<br>TD = -2.73, -2.76 |
| 8 | 43,575-43725 kb | *Hp*, TD | *LOC102183789* | - | -ZHp = 5.04, TD = -2.74 |
| 8 | 43,650-43,800 kb | *Hp*, TD | *KANK1* | Coat color | -ZHp = 5.21, TD = -2.77 |
| 8 | 45,450-45,600 kb | *Hp*, TD | *APBA1* | Body mass | -ZHp = 5.05, 5.08,<br>TD = -2.73, -2.73 |
| 12 | 33,750-33,900 kb | *Hp*, TD | *MYCBP2* | Body weight | -ZHp = 5.47, TD = -2.72 |
| 12 | 44,250-44,400 kb | *Hp*, TD | *LOC102173756* | - | -ZHp = 5.38, TD = -2.72 |
| 20 | 14,250-14,400 | *Hp*, TD | *ADAMTS6* | Body height | -ZHp = 5.63, TD = -2.74 |

## 4. Discussion

This genome wide selection scan presents the results of regions under positive selection using complementary statistical tests, – ZHp and TD on Kamori breed and also of genomic diversity analysis applied on Kamori vs. Bezoar using F_ST_ approach.

By assessing the SNVs in Kamori breed using – ZHp and TD, we observed selection footprints that are under the influence of both natural and artificial selection. Integration of these two approaches highlighted several concordant regions that are under selection (Table 3) which possibly include true positive selection signals with more confidence. These regions harbor genes related to body weight and height, coat color and milk production. Several regions harbor genes controlling body weight, immunity and body height traits are under genetic hitchhiking selection in Kamori such as on Chr. 2, 8, 13, 15 and 16. Among them, *RALY* gene has also been identified in Iranian Markhoz goat responsible for its black and brown coat color [23]. Similarly, *RAPH1* and *STIM1* is associated with body weight while *TPX2, NCAPG* etc. are related to body height in humans and in mice [9, 20]. Moreover, regions solely under old selection are also observed by *Hp* test that are located on Chr.3, 8, 13 and 18. Some examples include *ZFPM1* that is under selection in African sheep for milk yield [18]. The *ACSF3* gene is reported to be under selection in Korean cattle for meat production [19]. The *PIEZO1, CDH15, TDP2, EIF2S2, DUSP15* genes have previously been found under selection in humans for body height trait [20]. All the positive selection signatures found in Kamori adds support to the particular characteristics of this breed.

Our work also revealed clear genomic differences between the two goats under study which can be attributed to the regions varying majorly in body height, coat colors and milk protein among others. It has been shown that *STAG1, SIK3, SLC25A21, CCDC91, ZNF609, MGA, ATAD2B* and *TET1* are associated with body height in humans and in various livestock species [20] which differentiates the goats included here. For the *MIPOL1* and *IRF4* genes, studies have suggested their potential role in eumelanin pigmentation in chicken plumage and in human hair color respectively [20, 25] thus, they may explain the dark brown coat color of Kamori. The *TRMT1L* gene associated with chicken adaptation and survival in hot conditions [21] appeared under selection in Kamori which is likely due to its habitat in hot environment of Sindh province. One selective sweep harboring *ATP2C1,* a candidate gene for milk protein composition in Holstein bulls [17], appeared on a differentiated genomic region in this milch breed. None of the selection signals observed by F_ST_ above the threshold are consistent with the signals generated by – *ZHp* and TD, indicating that only divergent selection is acting on these regions.

## 5. Conclusion

With the aim to identify selection footprints in Pakistani Kamori goat breed, we applied three statistical tests; – ZHp, TD and F_ST_. A total of 98,574 autosomal positive as well as divergent selection signals were found that are likely associated with body mass, coat color, wool type, milk production, immunity, environmental adaptation, body height and reproduction. Our findings call for further investigation of Pakistani goat genome with the outcome that can support for selection and breeding programs for improved production and adaptive traits.

## Acknowledgements

Authors acknowledged Prof. Dr. Tosso Leeb’s great supports to accomplish this research endeavor, and we also acknowledge Dr. Safdar Ali Fazlani for being helpful in the Kamori sample collection.

## Availability of data and material

Relevant data is available in the manuscript including supplementary files and supporting information.

## Code Availability

F_ST,_ TD scripts are available publicly, while in-house ruby script was used for *Hp* statistics, Manhattan and other supplementary plots were plotted using Bioconductor qqMan package using R.

## Authors Contribution

Rashid Saif (RS) envisaged the idea, involved in critical thinking, analysis of data, editing, proofread and correspondence with journal. Jan Henkel (JH) was involved in software and Linux features understandings, analysis of data and provision of in-house *Hp* script. Tania Mahmood (TM) and Aniqa Ejaz (AE) helped in data analysis and initial write-up. Saeeda Zia (SZ) helped in understanding the statistical methods.

## Ethics Approval

All Pakistani animal were sampled with the consent of their owners by obeying the local regulations.

## Consent to participation

As this is not a human based study, but authors informed and took farmer’s consent before collecting samples of their animals.

## Consent for publication

Authors have taken permission from the PI of this SNSF funded project to further analyze this publicly available data and has no objection to publish this work.

## Supplementary Material

**Fig. (S1).** Illustration of quality checks results.

**Fig. (S2).** SNP density plot.

**Fig. (S3).** Distribution of – *ZHp* scores

**Fig. (S3).** Distribution of TD scores

**Fig. (S3).** Distribution of F_ST_ scores

**Table S1.** Genomic selection signatures −*ZHp* ≥ 5

**Table S2.** Genomic selection signatures TD ≤ −2.72

**Table S3.** Genomic selection signatures FST ≤ 0.09

