## Supplementary material for "Whole Genome Selective Sweeps Analysis in Pakistani Kamori Goat": Supplementary_Figures-KAM.pdf


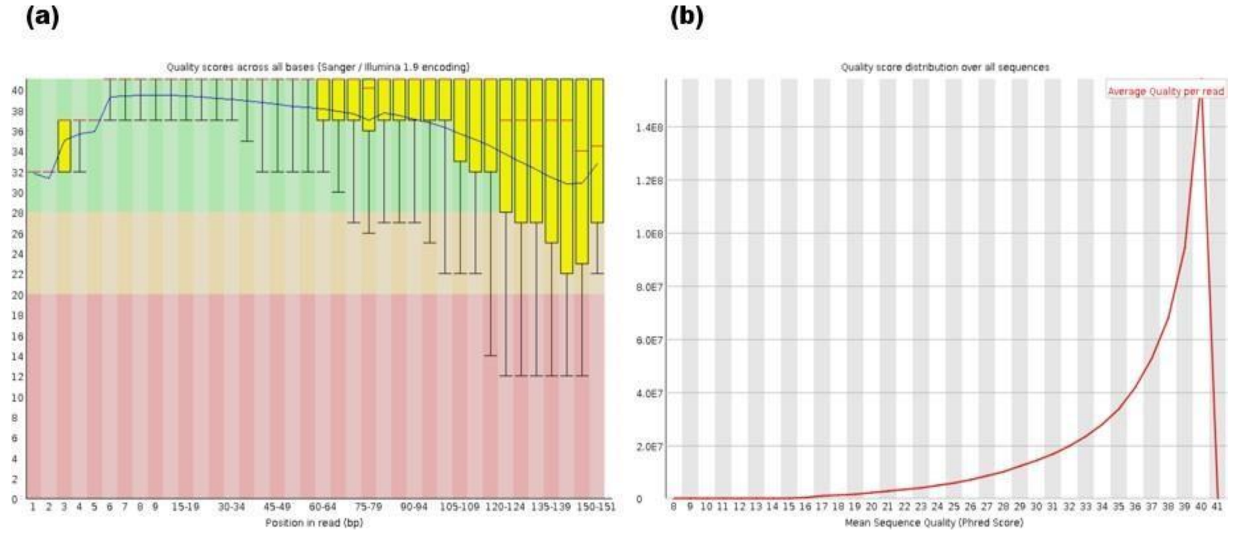


**Fig. S1** Illustration of quality checks results. (a): Quality score across all bases in bp is shown on x-axis vs. the average quality per read (phred scores) on y-axis. The yellow bar demonstrates interquartile range, red line indicates median value while blue line signifies mean quality of bases across the whole genome. (b): per sequence quality scores graph is representing phred scores on x-axis. The value above 27 indicates a high quality sequence score.

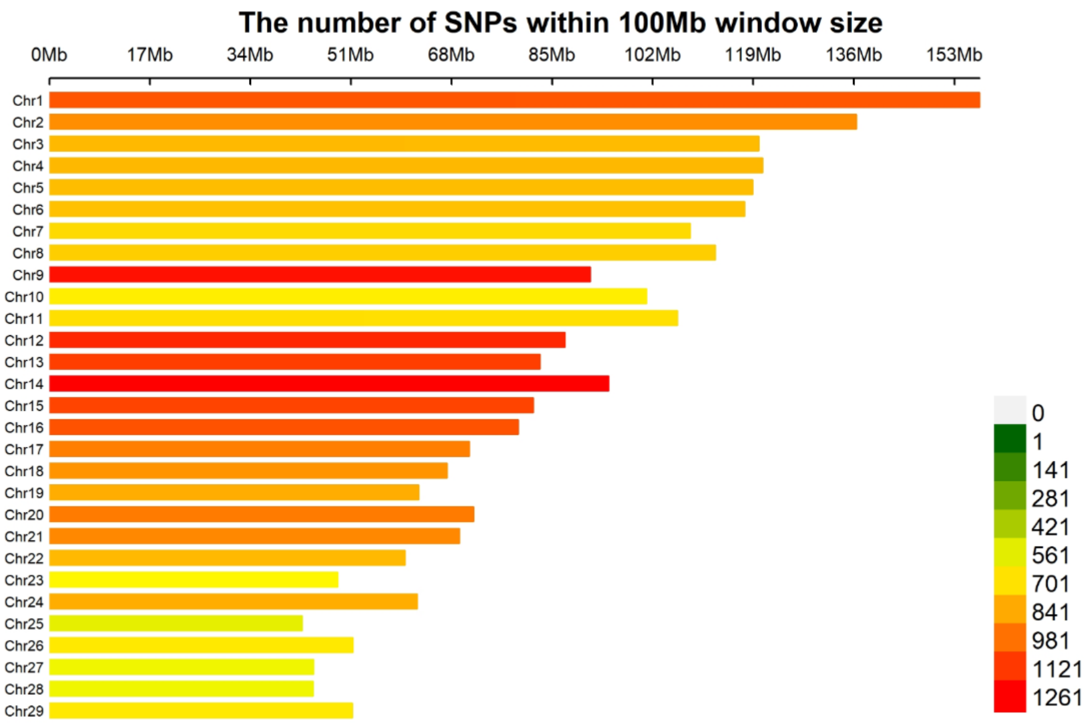


**Fig. S2** SNP density plot. The distribution of SNPs across all autosomes is illustrated. The number of SNPs within 100Mb window size is displayed on x-axis against the chromosome number on y-axis. The color gradient from low to high is represented as color key.

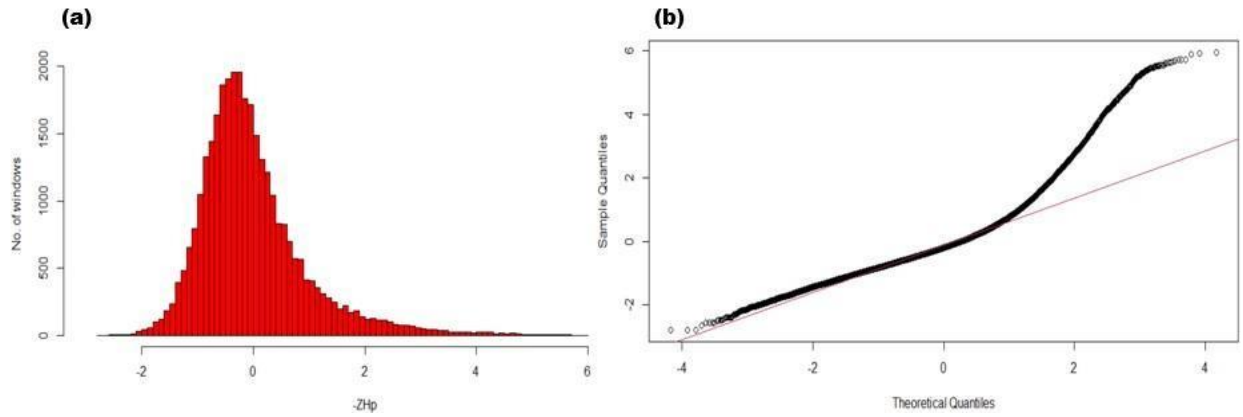


**Fig. S3** Distribution of $-\boldsymbol{ZHp}$ scores. (a): Frequency of windows under selection at a particular $-ZHp$ value is illustrated as histogram. (b): Q-Q plot of $-ZHp$ scores constructed shows the theoretical quantiles on x-axis vs. sample quantiles on y-axis. Red line in the plot indicates the expected distribution of SNP markers.

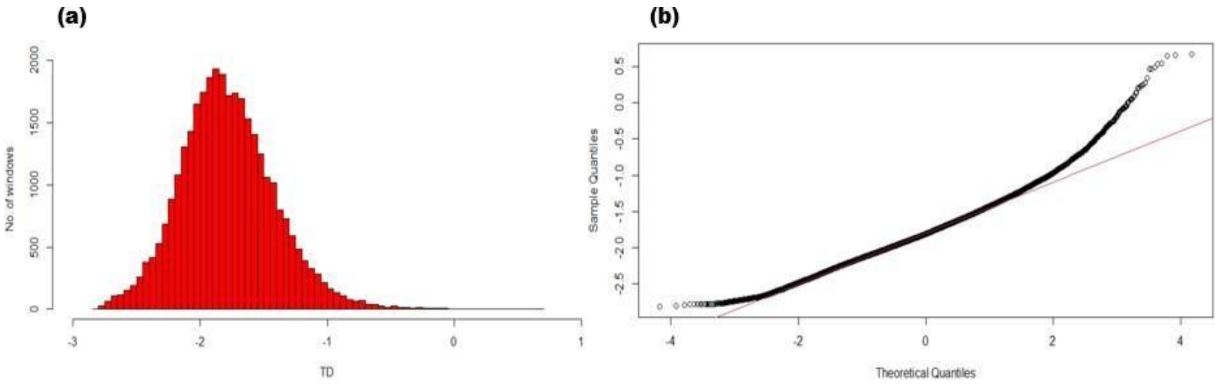


**Fig. S4** Distribution of TD values**.** (a): Frequency of recent selective sweep regions based on TD values are plotted on histogram. (b): Q-Q plot displays the comparison of observed distribution of TD values against theoretical model distribution of expected TD scores.

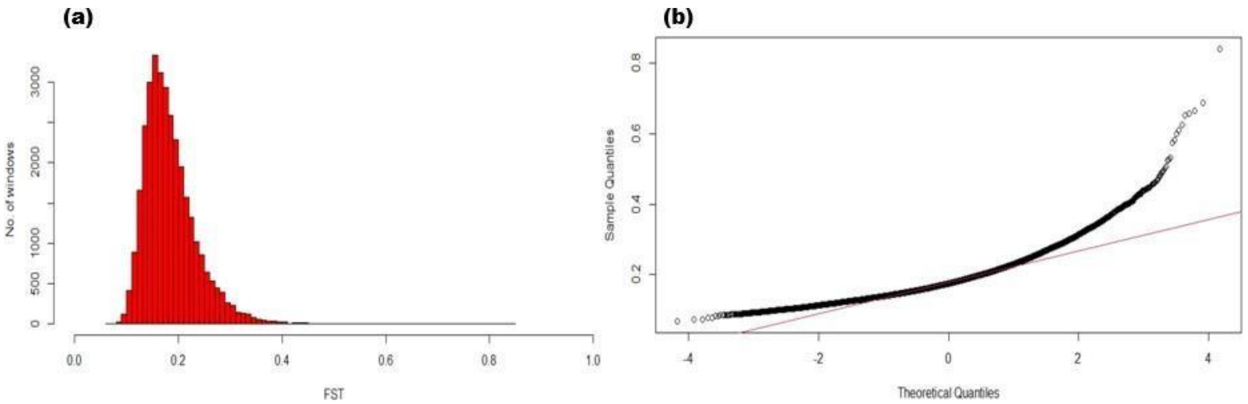


**Fig. S5** Distribution of $F_{ST}$ values**.** (a): The $F_{ST}$ values are plotted on x-axis vs. the number of times a particular $F_{ST}$ value occurred is displayed on y-axis as number of windows. (b): The sample quantiles in a Q-Q plot are plotted on y-axis against the expected quantiles of the standard normal distribution on x-axis. Red diagonal line indicates the theoretical distribution of $F_{ST}$ values across all autosomes while the black line is the observed quantiles.
